# Future state prediction errors guide active avoidance behavior by adult zebrafish

**DOI:** 10.1101/546440

**Authors:** Makio Torigoe, Tanvir Islam, Hisaya Kakinuma, Chi Chung Alan Fung, Takuya Isomura, Hideaki Shimazaki, Tazu Aoki, Tomoki Fukai, Hitoshi Okamoto

## Abstract

Human predicts future. To ask if fish also has this capacity, we established the virtual reality training system with live imaging of the telencephalic neurons of adult zebrafish in the active avoidance and found that, at the onset of the trial, learned fish conceives two future conditions as the favorable status on its way to the safe goal, *i.e.* one with the backwardly moving landscape and the other with the color of the safe goal. And the two different neural ensembles monitor the discrepancy between these predictions and the perceived real external status. Once fish reaches the goal, another ensemble is set to work to monitor whether fish keeps staying in the safe goal. The manipulation to artificially enhance these prediction errors elevated the activities of these ensembles and induced fish to behave to correct errors, revealing that fish sets behavioral strategy to actively realize these predictions.

## Introduction

Selecting optimal action in decision making according to the current sensory input is essential for animals. The optimal behavioral selection is thought to be taken based on the expected reward value and animals act to maximize its value (Sutton and Barto, 2018). Two strategies are proposed as the mechanisms; a model-free strategy in which animals directly attach the values to the actions or a model-based strategy in which animals make decisions based on the predictions by the internal model of the animal's brain which copies the structure of the external world (Daw et al., 2005). However, it is not yet clear how these two mechanisms work and interact in the actual brain. In this research, in order to reveal the mechanism of optimal behavioral selection, we established the closed-loop virtual reality 2-photon calcium imaging system for adult zebrafish. The zebrafish telencephalon has evolutionarily homologous regions and neural circuits to those of other vertebrates including mammals (Mueller and Wullimann, 2009), such as the cortico-basal ganglia circuit, which is implicated in behavior selection (Cui et al., 2013). Its brain is very small, allowing us to observe neuronal activities from a relatively wide brain region (Aoki et al., 2013). In addition, the use of the pigment deficient mutant strains enables observation of the telencephalic neuronal activities without opening the skull. These make adult zebrafish a novel attractive model animal to reveal the evolutionarily conserved and universal mechanisms of action selection in vertebrates.

Here we studied the active avoidance behavior as an example of the simplest behavioral paradigms to select optimal actions based on the value of the chosen environment, *i.e.* escape from the aversive to the neutral environment (Amo et al., 2014; Aoki et al., 2013; Dayan, 2012; Pradel et al., 1999). Although the active avoidance has been regarded as a typical example of the model-free decision making behavior, our results to our surprise revealed that the zebrafish telencephalon generates the predictions of future favorable states in its internal model and use the prediction errors from the external states to guide the fish to correctly reach the goal in the active avoidance.

Capacity to predict the desirable future and to plan the proper behavioral strategy to realize it has been regarded as the inherent functions of the higher vertebrate brain. However, our data show fish can do both.

## Results

### Establishment of the closed-loop virtual reality system for 2-photon real-time imaging of the telencephalic neuronal activity in behaving transparent adult zebrafish

To reveal the mechanism of optimal behavior selection, we established a closed-loop virtual reality 2-photon calcium imaging system. For this purpose, we first developed a technique to fix adult zebrafish head by attaching the custom-made harness with dental bond and cement (Figures 1A, S1A and S1B for detail see STAR Methods). Combined with the use of the transparent mutant strains (see STAR Methods), this method enabled continuous imaging of the beating tail and the telencephalic neuronal activities of adult zebrafish during trials under the virtual reality environment (Figures 1B, 1C and Movie S1) where the head-fixed live fish was put in a small tank surrounded by the four liquid crystal displays (LCD) on the left, right, front, and bottom of the tank. The presentation of the visual stimuli and the imaging of the neural activities were alternately performed by alternative switching of the LED backlight of the display and the photomultiplier tube (PMT) of the microscope (Figure 1D). During the scanning of each line (650.24 μsec), the Gallium arsenide phosphide (GaAsP)-PMT detector (Zeiss, BIG detector) was switched on and displays were switched off. During the shift to the next line (approximately 90 μsec), GaAsP-PMT was switched off and displays were switched on. To achieve this, we picked the TTL signal from the 2-photon microscope when scanning in line was started. The electronic stimulator (SEN_3401, NIHON KOHDEN) received this TTL signal and sent modified time TTL signal based on the original TTL signal derived from the microscope. This TTL signal from the stimulator was used to switch on and off the display. The switch of GaAsP-PMT was controlled by the custom-made system made by Zeiss.

**Figure 1.**
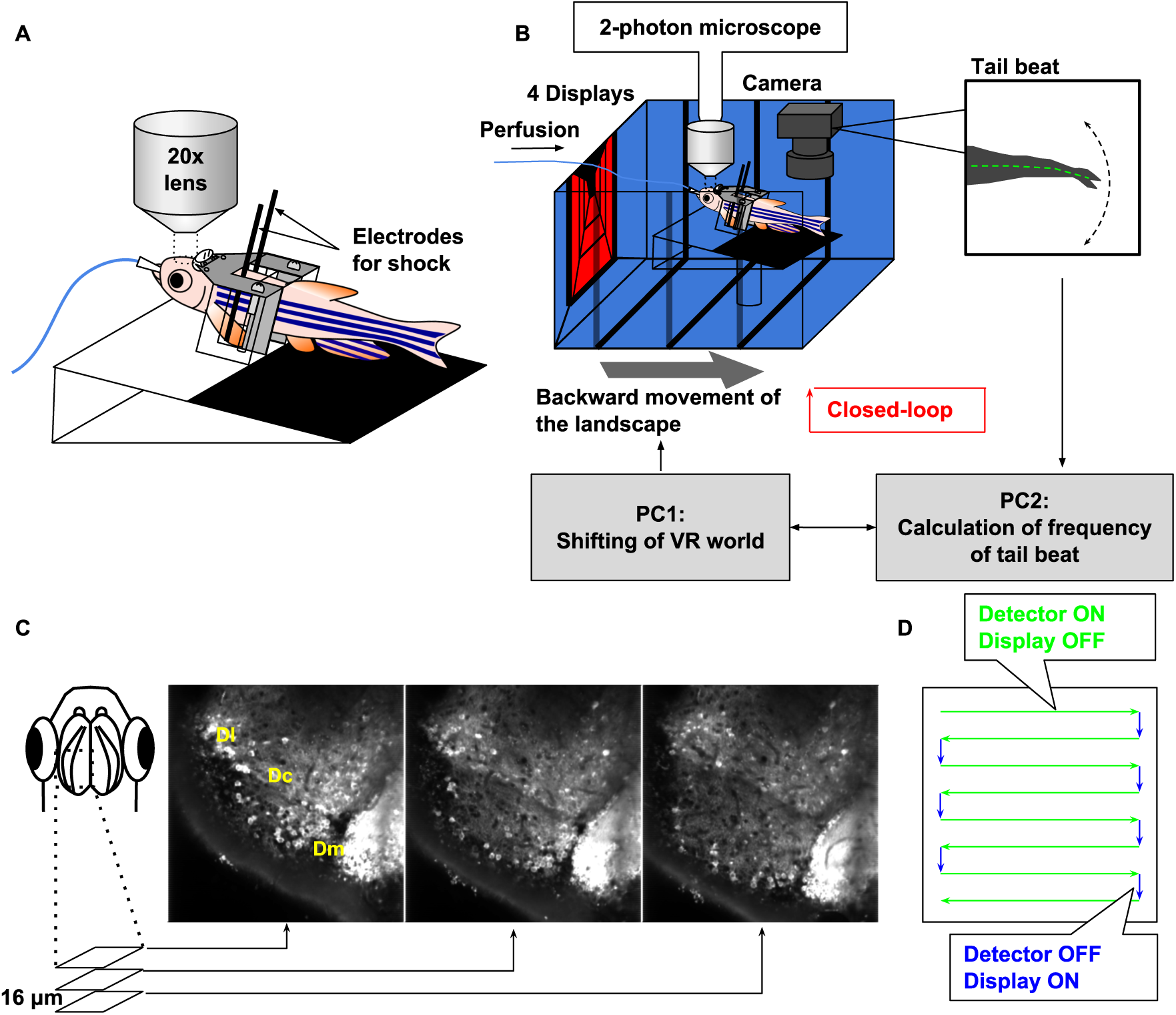
The closed-loop virtual reality 2-photon imaging system enables real-time capturing of the neuronal activities from behaving adult zebrafish. (A) Schematic drawing of the tethered living adult zebrafish using the custom-made harness, dental bond and cement (See also Figure S1.). The neuronal activities of the telencephalon were captured by 2-photon calcium imaging. Two needle electrodes were placed on both sides of the body of the tethered zebrafish to deliver electric shock. (B) Schematic diagram of the closed-loop virtual reality setup. Four displays presented visual stimuli. When fish beat the tail, landscape moved backward (gray arrow) to give fish the image of forward fictive swimming. The virtual traveling distance was calculated by [frequency of tail beats] x [gain]. (C) The calcium imaging of neural activities in three focal planes by using the piezo actuator. Either the left or right hemisphere was imaged. Anterior to the top; lateral to the left; medial to the right. Dl, lateral zone of dorsal telencephalic area; Dc, central zone of dorsal telencephalic area; Dm, medial zone of dorsal telencephalic area (Mueller and Wullimann, 2009). (D) Schema of alternative switching of neuronal activity detection by 2-photon microscope and visual stimulation by LCDs.

As fish beat the tail, the visual stimuli on the display were shifted backward according to the calculated traveling distance (Figure 1B, gray arrow) to make the system as the closed-loop (Figure 1B). We performed calcium imaging of neuronal activities in the telencephalon using the piezo actuator which allowed us to capture 3 slices of images separated by 16 μm (approximately 100 ms/slice)(Figure 1C and Movie S1). In the imaged fish, excitatory neurons were labeled with the genetically encoded calcium indicator G-CaMP7 (Ohkura et al., 2012) under the control of the camk2a and vglut2a promoter.

### Zebrafish can learn to take optimal actions to avoid shock under given rules in the virtual reality

To reveal the mechanisms for the adaptive goal-directed behavior, we designed the GO (active avoidance) / NOGO (passive avoidance) trials in the virtual reality (Figure 2A, upper panel). After the inter-trial interval (15seconds) where fish was surrounded with white background with black stripes, the GO or NOGO trials was randomly initiated. In the GO trial, the background color turned to blue and fish had to escape to the red region in their front within 10 seconds. For the NOGO trial, the background color turned to red and fish had to stay in the red region for 10 seconds. If fish did not behave as required, electric shock (5V/cm for 1second) was delivered from the two needle electrodes put on both sides of the body of zebrafish (Figure 1A). In some fish, after learning the original rule, the shock-associated color was reversed from blue to red, so that the fish had to escape to the blue region in the GO trial and stay in the blue region in the NOGO trial under the reversed rule (Figure 2A, lower panel).

**Figure 2.**
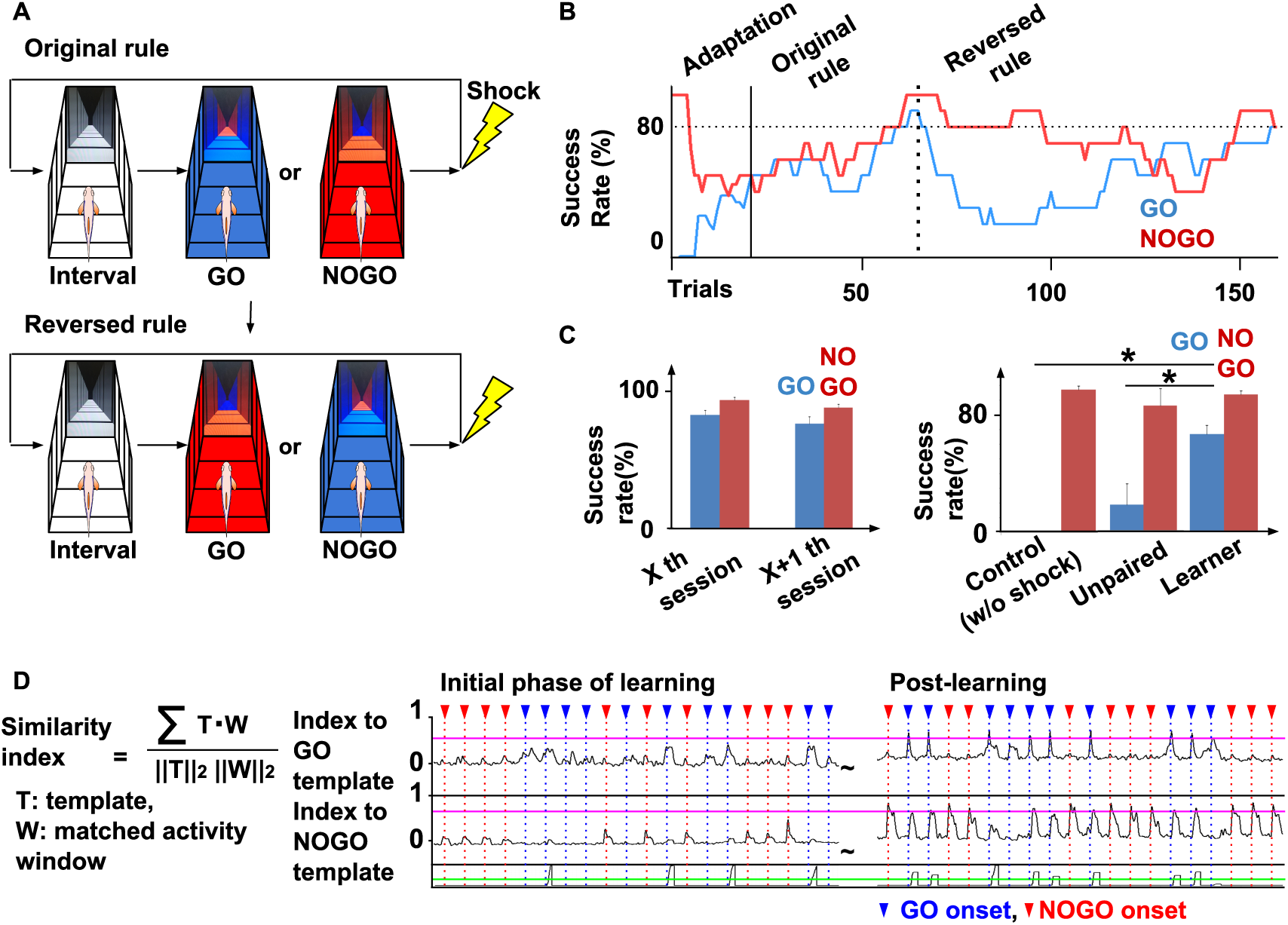
Fish can learn the GO/NOGO trials in the virtual reality and the specific neural population of the telencephalon is recruited during the correct avoidance behavior. (A) GO/NOGO trials in the original and the reversed rules. After the inter-trial interval (15 seconds), The GO or NOGO trial (10 seconds) was randomly initiated. In the original rule, electric shock (5V/cm, 1 second) was associated with the blue background color. In the reversed rule, shock was associated with the red background color. (B) Fish can learn the GO/NOGO trials in the original and the reversed rules. Learning curve of GO/NOGO trials in the original and the reversed rules. Success rate was calculated by the last 10 of each GO or NOGO trials. The blue and red lines indicate the results of GO and NOGO trials, respectively. 80% success rate was defined as the threshold for successful learning in both trials (horizontal dotted line). Vertical line, the initiation time point of trials by the original rule; Vertical dotted line, the timing of rule change. (C) Fish can learn and store the rule of the GO/NOGO trials. The left shows that the success rate of last 10 trials of GO and NOGO trials in the session fish satisfied the learning criteria (X th session) and that of first 10 trials of GO and NOGO trials in the next session (X+1 th session). There is no significant difference between the success rate between X th session and X+1th session, suggesting the stored performance of active avoidance behavior. The right graph shows that the success rate in 29th GO trials and 12th NOGO trials in the control (without shock, n = 4), the unpaired (electric shock was delivered in the inter-trial interval, n = 4) and the learner (n = 13) (right). 29 and 12 were the average of the trial numbers to achieve the learner criteria in the GO and NOGO trials, respectively. Fish under the control or the unpaired condition did not show the increase of the success rate in the GO trials, suggesting no learning. *p<0.01, t-test. (D) Template matching analysis. The similarity between the template and matched activities in all time windows is calculated. Two templates were made averaging neuronal activities derived from success GO and success NOGO trials at the time when fish reached the learning criteria (see STAR Methods). The similarity showed an increase after learning, suggesting that the specific neuronal population was recruited for success in each trial category (n = 12). Blue and red triangles indicate the onset of GO and NOGO trials, respectively. Magenta line indicates mean + 3SD of similarity index. Black line in the lower graphs shows the distance that fish has traveled in the virtual reality space. Green line indicates the position of the goal.

Figure 2B shows the learning curve of the GO/NOGO trials under the original and reversed rules. During the adaptation period without shock, fish has no preference to the color (Figure 2B). However, as learning proceeded, fish gradually mastered the GO trial and eventually, the success rate of both GO and NOGO trials reached the learning criteria, *i.e.* 80%, in both trials, calculated from the past 10 trials (Figure 2B).

In some fish, after the original learning was achieved, we suddenly reversed the rule Figure 2B, vertical dotted line). The others continued the trials under the original rule in the following sessions. Fish were able to learn even after changing the rules (n =4). In the fish which continued the trials under the original rule, the success rate of the last 10 trials of GO (83.1 ± 3.28%) and NOGO (93.8 ± 2.13%) trials in the session in which fish achieved the learning criteria was not significantly different from that of the first 10 trials of GO (76.92 ± 5.23%) and NOGO (88.46 ± 2.74%) trials in the next session (GO trials, p = 0.312; NOGO trials, p = 0.089, t-test, n = 13) (Figure 2C, left panel). Compared with the control (without shock), and the unpaired fish, the fish which underwent learning trials had the higher rate of success (Figure 2C, right panel). These results show that fish could learn the GO/NOGO trials, adapt to the reversal of the rule and that fish learn to take the optimal actions to avoid shock under given rules even in the virtual reality.

### Specific neural populations are recruited for the optimal behavior in the learner fish

To reveal the neuronal basis of optimal behavioral selection, we analyzed data of calcium imaging of the telencephalic neuronal activities. To ask whether specific neuronal populations were recruited for the success trial, we first performed template matching analysis. For this analysis, we made the template using the neuronal activities data within 2 seconds from the onset of the trial either in the success GO trials and NOGO trials after learning. We calculated the similarity with the template (Figure 2D) at each time point by sliding the template from the onset to the last of the session (60 trials). As a result, when the GO trial template was used, the similarity at the onset of the GO trial became high after the learning (Figure 2D). We found the similar results in the case using the NOGO template. When the NOGO template was used, the similarity to the template showed a higher value at the onset of the NOGO trial after learning. These results demonstrate that specific neuronal populations were recruited for success in the GO and NOGO trials after learning. Taken together, by learning, specific population neurons were recruited for the optimal behavior.

### The NMF analysis of neural activities identified two neural ensembles encoding color perception and action while perceiving color

To further elucidate the nature of the specific neuronal populations, we performed Non-negative Matrix Factorization (NMF) (Lee and Seung, 2001) (Figure 3A). The timeline of neuronal activities can be expressed by one matrix with each row corresponding to the time-lapse activity of each neuron. The NMF factorizes this matrix into two with the columns of the first matrix showing the typical ensembles of neural activities and the row of the second one showing how frequently each ensemble shows up at each time point. The number of ensembles was determined according to Akaike Information Criteria (AIC) (Figure 3A). By comparing the activity timeline with the behavior data, we identified what kind of information is encoded in each neural ensemble. Throughout this paper, we use the term "activity" to refer to the emergence frequency of each ensemble.

**Figure 3.**
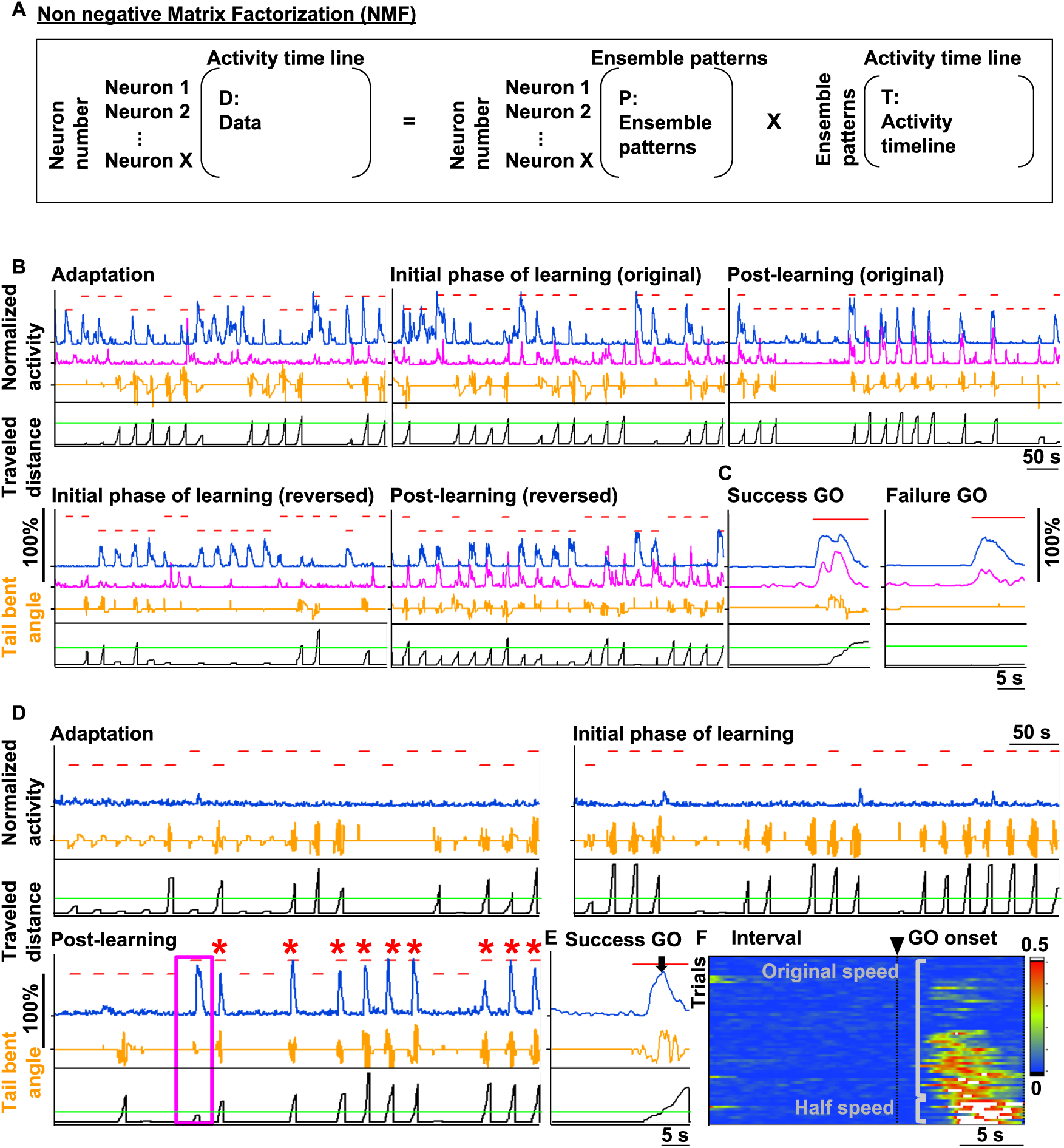
Non-negative matrix factorization reveals the neural ensembles showing different responses to the visual sensory input, movement and position of fish. (A) The formula to calculate the non-negative matrix factorization (for details see STAR Methods). (B) The activities of two neural ensembles in the original and reversed GO/NOGO trial. Blue and magenta lines indicate the activity of each neural ensemble. Blue ensemble showed up when fish perceived blue color regardless of learn or not. Magenta ensemble showed up when fish moved perceiving blue color after learning (upper right panel). The upper graphs indicate the activity of neural ensemble normalized by self-maximum value. In the upper graphs, red horizontal bars in the upper position indicate the GO trial period and those in the lower position indicate the NOGO trial period. Orange line indicates the tail bent angle. Black line in the lower graphs shows the distance that fish has traveled in the virtual reality space. Green line indicates the position of the goal. (C) The close-up view of a single success GO trial in the original rule (left panel) and a single failure GO trial in the original rule (right panel) in (B). Blue ensemble showed up when a GO trial started regardless of success or failure of the trial, and magenta ensemble showed up only when fish swam successfully. (D) The neural ensemble encoding the position in relation to the goal. The activity of a neural ensemble during the adaptation (upper left panel), the initial phase of learning (upper right panel) and after learning (lower panel). This ensemble showed up after learning and had a sharp peak around the goal in the success GO trial (asterisks). Interestingly, the width of the peak became thicker in some peculiar trials of the GO trial when fish stopped on the halfway to the goal (magenta boxed area). (E) The close-up view of a single success GO trial in (D). The activity of the ensemble shown in (D) in a single success GO trial after learning. Arrow indicates the time when fish reached the goal. This ensemble had a peak around the goal border (See also Figure S2.). (F) The appearance of the activity peak was delayed when the speed was halved by changing the "gain" constant from 10 to 5. The activity of the neural ensemble shown in (D) from all success GO trials are aligned. Arrowhead indicates the onset of the GO trial.

In Figure 3B, the ensemble shown in blue line was already active during the adaptation and the GO trial from the initial phase of the learning (upper left and middle panels). Its activity can be seen even after the establishment of learning (right panel). This ensemble showed up regardless of achievement of learning and the activities were also observed in the NOGO trial in the reversal learning where the same blue background was shown as in the GO trial in the original rule. Therefore, we interpreted this ensemble as coding perception of blue visual stimuli because its activity correlated very well with whether the fish was perceiving the blue background color (Figures 3B and 3C, blue line). On the other hand, the ensemble shown in the magenta line showed up when fish was swimming forward correctly in the GO trial of the original learning and mistakenly swimming in the NOGO trial of the reverse learning (Figures 3B and 3C, magenta line), suggesting that it showed up when fish was swimming forward while receiving the blue visual stimulus. Thus, we identified one ensemble encoding blue color perception and another that encodes action (swimming) while perceiving the blue color background (n = 3).

### The neural ensemble encoding the position in relation to the goal

We also identified another ensemble which emerged in relation to the goal. This neural ensemble hardly showed up during the adaptation (Figure 3D, upper left panel) nor the initial phase of learning (Figure 3D, upper right panel). This ensemble showed up when fish succeeded in the GO trial after learning (Figure 3D, bottom left panel). Interestingly, its time-lapse pattern was steep in the success GO trials (Figure 3D, asterisks in the bottom left panel) but became blunt when fish stopped halfway to the goal (Figure 3D, magenta boxed area). From these results, we speculate that this ensemble encodes the position in relation to the goal with a peak around the goal border (Figure 3E, arrow) (n = 6). To test this possibility, we performed speed change experiments by reducing the gain of the tail beat, with the prediction that when the speed is halved, the appearance of the peak of the activity should be delayed because fish reaches the goal later. In fact, in the case of reduced gain, the peak of the activity of the ensemble shifted behind as expected (Figure 3F). Average time to reach the peak after initiation of the GO trial shifted from 5.64 ± 0.44 seconds under the original gain (last 9 success GO trials) to 6.91 ± 0.53 seconds after halving the gain (9 success GO trials) (p = 0.043, t-test). In addition, the constituent neurons of this ensemble retained the sequential order of activation throughout all 10 success GO trials (Figure S2), suggesting the robustness of position encoding by the neurons in this ensemble.

### The elevation of the activity of the ensemble encoding the error with respect to the prediction of the goal color induces tail beats

Intriguingly, unlike the neural ensemble that encodes the perception of blue color, we identified the other neural ensemble whose activity increased specifically after learning. This ensemble did not show up during the adaptation (Figure 4A, upper left, blue line) nor the initial phase of learning (Figure 4A, upper right, blue line), but showed up only after learning (Figure 4A, lower panel, blue line), suggesting that this ensemble shown in the blue line does not merely represent the perception of blue color. Focusing on one trial, this ensemble showed a decrease in activity as fish was reaching the goal (Figure 4B, blue line). In other words, this ensemble showed a decrease when fish perceived a sensory input of red color as a favorable goal color. We also observed another ensemble which complementarily increased activity when fish reached the red goal region (Figure 4B, magenta line). These results suggested that the neural ensemble shown in the blue line encodes an error with respect to the prediction of perception of the red background color as a favorable goal color.

**Figure 4.**
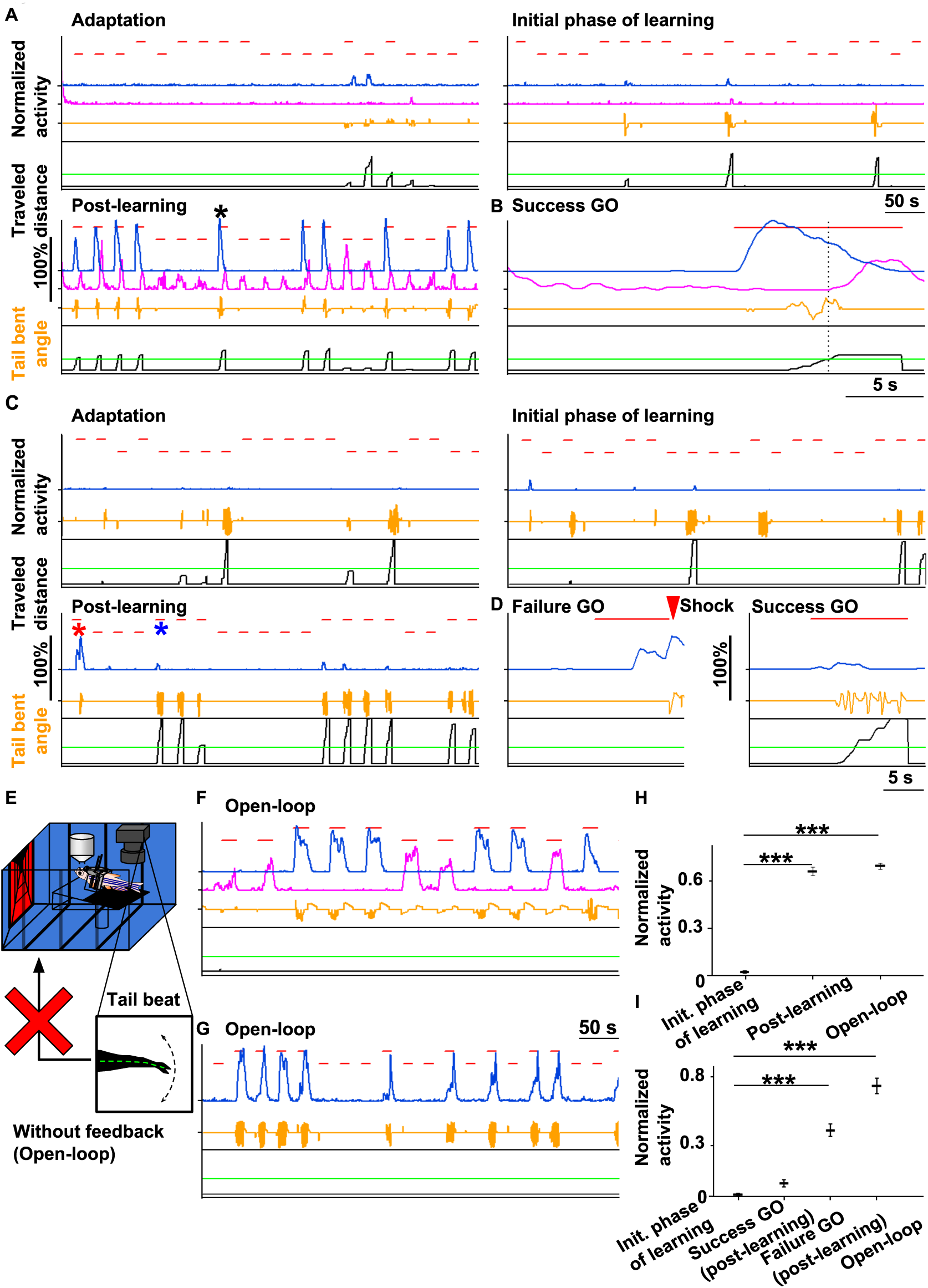
Open-loop experiment in the GO trial reveals the neural ensemble which encodes the errors between predicted favorable states and the perceived states. Notations in the figures are all same as in Figure 3. (A) The activity of two neural ensembles during the adaptation (upper left panel), the initial phase of learning (upper right panel) and after learning (lower panel). Both of these ensembles showed up in the GO trial as learning proceeded but their onsets were different timing. (B) The close-up views of the activity of the two ensembles shown in a single success GO trial after the establishment of learning indicated by an asterisk in (A). Vertical dotted line indicates the time point when fish reached the goal. The blue ensemble showed up when fish was presented with the blue background, and subdued as fish approach the goal. The magenta ensemble showed up after passing into the goal with the red background. (C) The activity of a neural ensemble during the adaptation (upper left panel), the initial phase of learning (upper right panel) and after learning (lower panel). This ensemble showed up only when fish failed in the GO trial after learning (red asterisk). (D) The close-up views of the activity of the neural ensemble as indicated by red and blue asterisks in (C) in a single failure GO trial (left panel) and a single success GO trial (right panel) after learning. Red arrowhead indicates the onset of electric shock for punishment. (E) The open-loop experiment, where the feedback was turned off. The landscape did not move in response to the tail beat. (F) The activity of two ensembles shown in (A) in the open-loop condition. The blue line ensemble showed up and kept elevated by the end of the GO trial. Fish kept beating the tail during the GO trial as shown by orange line. The magenta line ensemble did not show up throughout this trial. (G) The activity of the neural ensemble shown in (C) in the open-loop condition. The ensemble showed up and kept elevated by the end of the GO trial in contrast to the closed-loop condition, where this ensemble did not show up at all (C). Fish kept beating the tail during the GO trial as shown by orange line. (H) Comparison of the peak activity of the ensemble shown in the blue line in (A), (B) and (F) (see STAR Methods). Error bars: mean ± SEM. ***p<0.001, one-way ANOVA, Bonferroni's multiple comparison test. Initial phase of learning, 0.0268 ± 0.0086; Post-learning, 0.6501 ± 0.0253; Open-loop, 0.6832 ± 0.0181. Calculated by the peak values from GO trials in each different phase of learning and open-loop condition. (I) Comparison of the peak activity of the ensemble shown in the blue line in (C), (D) and (G) (see STAR Methods). Error bars: mean ± SEM. ***p<0.001, one-way ANOVA, Bonferroni's multiple comparison test. Initial phase of learning, 0.0134 ± 0.0080; Success GO, 0.0797 ± 0.020; Failure GO, 0.3892 ± 0.0361; Open-loop, 0.6501 ± 0.0457. Calculated by the peak values from GO trials in each different phase of learning and open-loop condition.

To confirm that this ensemble encodes a prediction error, we artificially manipulated the experimental condition so that the error in this respect would stay elevated by putting fish in an open-loop environment in which fish could not swim forward however hard fish beats the tail in the GO trial. In the open-loop, indeed, the activity of this ensemble was sustained during the trial (Figures 4F and 4H), and fish kept beating the tail during the trial (Figure 4F, orange line). Therefore, this neural ensemble encodes the error between the prediction of perception of the red goal color and the real visual input. Besides, we observed that fish kept beating the tail throughout the open-loop trial. This observation demonstrated that the elevation of this error induces tail beats, and fish keeps beating the tail unless this error signal disappears (n = 9).

### The elevation of the activity of the ensemble encoding the error with respect to the prediction of visual perception of backward movement of landscape induces tail beats

Furthermore, we also observed another neural ensemble which increased activity very prominently in the failure GO trial after learning (Figures 4C and 4D). During the adaptation (Figure 4C, upper left panel), the initial phase of learning (Figure 4C, upper right panel) and the success GO trials after learning (Figure 4D, right panel) its activity was not observed. Its activity increased only in the failure cases of the GO trial (Figure 4D, left panel, red asterisk). Therefore, this ensemble showed up only when fish did not beat the tail and subsequently did not perceive the backward movement of the landscape (Figures 4D and 4I). Under the open-loop condition where the error with this respect remains to be elevated as the landscape stays still irrespective of the fish tail beat, the activity of this ensemble was sustained to be elevated (Figures 4G and 4I), and fish continued vigorous beating of the tail throughout the trial (Figure 4G, orange line), demonstrating that this ensemble encodes another error between the prediction of visual perception of backward movement of the landscape and the real visual perception, and fish keeps beating the tail unless this error signal disappears (n = 6).

### Another neural ensemble was identified which encodes the error with respect to the prediction of the goal color after reaching the goal

We further wondered whether fish has any prediction and detect prediction error even after fish reached the goal. To examine this possibility, the color was changed to green or white as soon as fish reached the goal (Figure 5A). Interestingly, we found a new ensemble which showed up specifically with the change of the goal color in the GO trials (Figure 5B). This neural ensemble showed up regardless of the changed color of the goal (Figures 5B, green and open circles, 5C and 5D), suggesting that this neural ensemble encodes an error between the perception of the expected goal color where fish has to stay and the real visual input after reaching the goal (n = 3).

**Figure 5.**
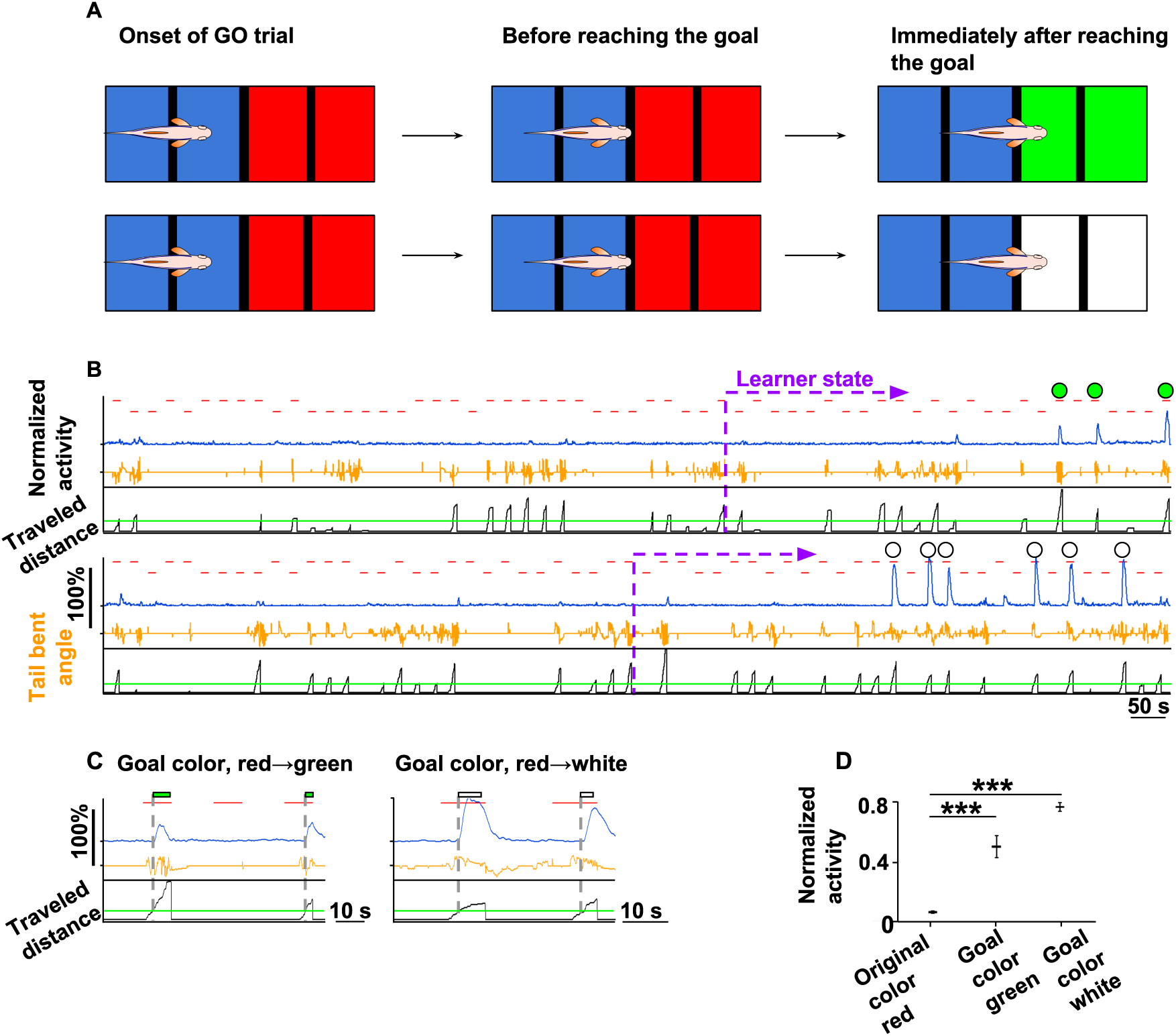
Goal color change experiment reveals the ensemble encoding the error between actual perceived goal color and expected red goal color. (A) Schema of the goal color change experiment. In the GO trials, we changed the background color of the goal from red to green or white as soon as fish reached the goal. (B) The activity of a new neural ensemble. Green circles indicate the GO trials in which the goal color was changed to green. White circles indicate the GO trials in which the goal color was changed to white. This ensemble showed up in the GO trial in which the goal color was changed either to green or white. (C) The close-up views of the activity of a neural ensemble shown in (B). left panel, the two success GO trials in which the goal color was changed to green; right panel, the two success GO trials in which the goal color was changed to white. The dotted line indicates the time point when fish reached the goal and the goal background was changed. The changed colors were indicated by rectangles in corresponding colors. (D) The comparison of the peak activity of the ensemble depending on the goal color. The ensemble showed up only when the goal color was changed either to green or white from the original red color. Error bars: mean ± SEM. ***p<0.001, one-way ANOVA, Bonferroni's multiple comparison test. Original goal color red, 0.0389 ± 0.0047; Goal color green, 0.4598 ± 0.0842; Goal color white, 0.8103 ± 0.0411. Calculated by the peak value from success GO trials in each goal color.

## Discussion

Taken altogether, our results demonstrated that fish generates predictions by learning on three different aspects of the sensory status as the intermediate goals in order to successfully carry out the GO trial (active avoidance), and the prediction errors represented by different neural ensembles are monitored, and behaviors are taken so that these errors are minimized during the trial as summarized below (Figure 6).

**Figure 6.**
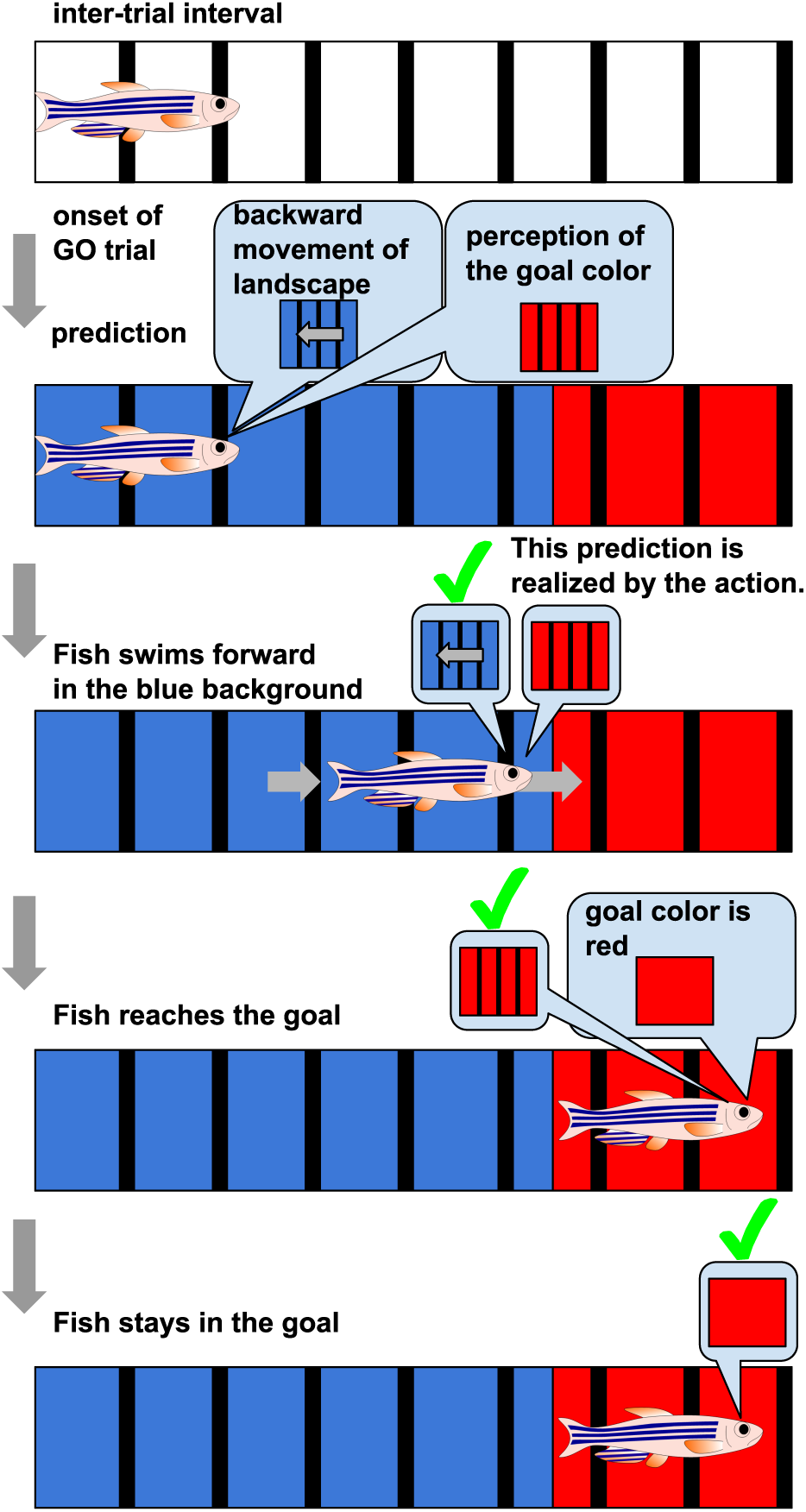
Realization (indicated by check marks) of predictions for the fulfillment of the active avoidance behavior.

First, during the inter-trial interval when fish perceives the white background color, there is no prediction, and there is no prediction error. Second, when the GO trial starts, fish perceives the blue background color and makes two predictions, *i.e.* visual perception of backward movement of the landscape and perception of red as the favorable goal color. Third, by selecting the optimal action, *i.e.* swimming forward, the error from the first prediction disappears. Fourth, when reaching the goal and perceiving red as the favorable goal color, the second error also disappears. Simultaneously, another prediction emerges that designates fish to continue to see the red goal color once reaching the goal. Thus, the correct behaviors of the learned fish, *i.e.* swim in blue and stop in red are carried out so that the errors with respect to these three predictions are minimized.

The prediction error coding ensembles were formed on the basis of past successful experiences. Actually, we identified multiple neural ensembles whose activity showed up only after learning and did not disappear unless the fish receives the optimum visual inputs as predicted. These ensemble activities stayed elevated in the open-loop environment in which errors could not be reduced by any behavior, further confirming that these neural ensembles encode prediction errors. And the continuous intense swimming observed during the open-loop trials demonstrated that fish chose the behaviors to minimize these errors as much as possible.

The active avoidance has been regarded as the most typical model-free decision making behavior (Amo et al., 2014; Aoki et al., 2013; Dayan, 2012). However, our results show that it proceeds with monitoring the errors from the predictions which are generated by the internal model, making the theoretical distinction between the model-free and model-based decision making behaviors obscure.

Our results are in good agreement with the data from other groups that the zebrafish brain is capable of the predictive coding as those of other higher vertebrates (Attinger et al., 2017; Chao et al., 2018; Funamizu et al., 2016; Makino and Komiyama, 2015; Rao and Ballard, 1999; Schneider et al., 2018; Schwiedrzik and Freiwald, 2017; Sumbre et al., 2008).

Our results are also in line with the behavioral control scheme so-called “active inference” where actions are selected so that they will minimize the discrepancy from the model-based sensory predictions by altering the sensory environments by action (Pezzulo et al., 2015; Dayan et al., 1995; Friston et al., 2017).

Some groups have reported the telencephalic neurons both in mouse and adult zebrafish with the similar response as the neural ensemble which we have identified here to be encoding the error from the prediction of the backward movement of the landscape (Attinger et al., 2017; Huang et al., 2019). Although both populations are activated under the similar situations where the animal actions cause unexpected movement of the landscape, which includes the case of no movement under the open-loop condition, the ensemble reported here is different from others in that it was also activated when fish failed to correctly beat the tail in response to the conditioned stimulus and no movement of the landscape was induced. Therefore, the ensemble we have reported here is not the same type of neural population which simply encodes the error from the prediction of the landscape movement in response to the animal’s action.

Previously, we found that the ventral habenula neurons encode the expectation value of the incoming punishment (Amo et al., 2014). In the context of active avoidance, these neurons show a gradual increase of tonic activation in response to the conditioned stimulus at the initial stage of association between the conditioned and unconditioned stimuli. However, as the learning proceeded, the level of activation of these neurons in response to the conditioned stimulus gradually returns eventually back to the basal level. In contrast, the neural ensemble which encodes the error from the prediction of the favorable goal color continues to be activated by the conditioned stimulus even after the establishment of learning. Therefore, the neural ensemble we identified has the completely different nature from that of the previously reported neurons in the ventral habenula.

In this study, we did not find the neural ensemble which directly encodes the predictions of sensory perception. In the predictive coding, the top-down predictions are expected to act inhibitorily to cancel the bottom-up sensory inputs for the derivation of the prediction errors (Makino and Komiyama, 2015; Schneider et al., 2018; Schwiedrzik and Freiwald, 2017), suggesting that the predictions may be represented by the GABAergic neurons in the telencephalon. By using other transgenic line expressing G-CaMP7 in the inhibitory neurons, we may be able to examine this possibility.

Because of its evolutionary conservation in its basic structure, the zebrafish brain can now be regarded as the miniature of the mammalian brain (Aoki et al., 2013; Mueller and Wullimann, 2009). In fact, it has the evolutionarily homologous structure to the isocortex, the hippocampus, and the cortical amygdala in its pallium and to the striatum and the globus pallidum in its subpallium (Aoki et al., 2013; Lal et al., 2018; von Trotha et al., 2014; Wullimann, 2014). In addition, the zebrafish brain has also the neuromodulatory systems such as those of dopamine and serotonin as in the mammalian brain (Amo et al., 2014; Lillesaar et al., 2009; Rink and Wullimann, 2002; Yamamoto et al., 2010, 2011). All these structures are known to play important roles in decision making behaviors. The small size of the zebrafish brain enables the observation and manipulation of much wider regions of the live brain than by using the mammalian brain. Therefore, the adult zebrafish brain can be an excellent model to elucidate how neurons in these homologous elements of the brain function and interact at the neural circuits and subcellular levels to generate the internal model for prediction and to use it for behavioral control in decision making.

## Acknowledgment

We thank Drs. Junichi Nakai, Tetsuya Koide and Yoshihiro Yoshihara for providing G-CaMP7 plasmids, Dr. Masae Kinoshita for drawing the excellent schema of adult zebrafish, Dr. Yukiko Goda for critical reading of the manuscript, Drs. Yuki Tanimoto and Ryo Aoki for technical assistance and discussion and Drs. Kuo-Hua Huang and Rainer W. Friedrich for communication before publication. We thank all members of the Okamoto laboratory for support and advice, Advanced Manufacturing Support Team of RIKEN for fabricating finely tuned apparatus for fish fixation, the Research Resource Center of RIKEN Center for Brain Science for fish care, Drs. Shinichi Higashijima and Chie Satou and the National BioResource Project of Japan for providing *TgBAC(vglut2a: Gal4)* fish. All materials, original data and custom codes for analysis are available upon request to H.O. and M.T. This work was supported by the RIKEN CBS Internal Budget, Strategic Research Program for Brain Science (H.O.) from MEXT and AMED, and Grant-in-Aid for Innovative Area (H.O., 23120008) from MEXT, the Core Research for Evolutional Science and Technology from JST and AMED (H.O., JPMJCR09S1), Grant from Kao Corporation (H.O.), Grant from Fujitsu Corporation (H.O.), the RIKEN Special Postdoctoral Researchers Program (M.T.) and Grant-in-Aid for Young Scientists (B) (M.T., 18K14858) from JSPS.

## Author contributions

H.O. conceived and led the project and M.T. and H.O. designed the experiments and analyses. T.Islam and M.T. built the virtual reality and computerized control and data acquisition systems. M.T. carried out the experiments. M.T., T.Isomura, H.S. and T.A. performed the analyses. C.C.A.F. and T.F. made the NMF program. H.K. generated fish line. M.T. and H.O. wrote the manuscript.

## STAR Methods

### Contact for Reagent and Resource Sharing

Further information and requests for resources and reagents should be directed to the corresponding author, Hitoshi Okamoto or Makio Torigoe.

### Animals

All surgical and experimental procedures were reviewed and approved by the Animal Care and Use Committees of the RIKEN Center for Brain Science. Zebrafish (*Danio rerio*) were bred and raised under the standard conditions (Weisterfield, 2007). In this study, we used *TgBAC(camk2a: GAL4VP16)^rw0154a^; TgBAC(vglut2a: Gal4)* (Satou et al., 2013)*; Tg(UAS: G-CaMP7) ^rw0155^* zebrafish more than 6 month old in the nacre (Lister et al., 1999)) or casper (White et al., 2008) background. We used *camk2a* containing BAC clones zH278O8 to establish *TgBAC(camk2a: GAL4VP16)* as shown in the previous paper (Amo et al., 2014) and *UAS: G-CaMP7* (Ohkura et al., 2012) plasmid was provided by Dr. Koide (Yabuki et al., 2016).

### Virtual Reality

The virtual reality environment consists of 4 liquid crystal displays (LCDs) (Good display, Model No. GD70MLXD) which present the visual stimuli. In this environment, fish can swim and stop along one dimensional track consisting of white, blue or red background color with black stripes at the right angle to the direction of swimming. The tail movement was captured by a web camera (BSW20KM11BK, iBUFFALO) with modification as follows. The filter was replaced with the infrared sharp cut filter (IR 76 FUJI FILTER, FUJIFILM) to avoid the interference from the display light. The tail was illuminated with infrared LED light (850 nm)(ILR-IO16-85NL-SC201-WIR200, intelligent LED solutions). The virtual reality system was controlled by custom-written programs based on LabVIEW (National Instruments), Matlab (Mathworks) and OMEGA SPACE (Solidray Co Ltd). The virtual reality environment was created by using OMEGA SPACE software, which provides built in tools for the creation of virtual space. As fish was fixated inside the tank, it could move its tail, and its tail beat frequency was continuously detected to cause forward-only motion in the virtual space. To investigate the frequency, the following was implemented. With a camera from top, and a custom program based on Matlab Computer Vision Toolbox and LabVIEW, the body of fish was detected. The camera was placed in such a way that the body of fish was along the vertical (Y) axis of the image taken by the camera. For each time frame of 100 ms, a centerline lying along the fish body was calculated by connecting the midpoints of the fish body in every row of the image. The upper points of the centerline corresponded to the upper portion of the fish body. The upper most portion of the fish body in view did not move during tail-bending, so we took 5 upper most points of the centerline and calculated a straight-line, which was at 90 degrees of inclination with the horizontal axis of the image. Next, we took 5 lower-most points in the centerline and fitted a straight-line comprising these. When fish bent its tail, this second line created an angle of inclination with the initial straight-line, which had 90 degrees of angle. When fish did not move its tail, this angle was 180 degrees, and the angle reduced when the fish moved its tail. So we subtracted this angle from 180 degrees to find how much the tail shifted from the reference position. This subtracted value was used as the tail beat angle. Furthermore, we observed that there was bias in direction of the tail angle depending on fish, so we took a comparatively large moving time window of 10 seconds and averaged the tail beat angles in that window to set the reference angle of the tail. In each time step, the difference between the reference and the tail beat angle was calculated to obtain the actual tail bend angle, which was used for calculation of tail beat frequency. To calculate the frequency, we performed Fourier transform using the current and 9 previous values of the tail beat angle, corresponding to 1 second of length in our system. The calculated frequency was multiplied by arbitrary constant "gain" to use as the forward speed of fish in virtual reality space, as the speed of fish was considered to have a linear correlation with tail beat frequency (Hunter R. and Zweifel, 1971).

### Fixation of living adult zebrafish

To capture the tail movement and image the telencephalic neural activities, fish were tethered to the custom-made harness. Adult zebrafish were briefly anesthetized with 0.02% tricaine (ethyl 3-aminobenzoate methanesulfonate salt, Sigma-Aldrich) diluted in fish rearing water and mounted in a hand-made surgery apparatus. During surgery, fish rearing water with 0.02% tricaine was continuously perfused to keep fish alive and its anesthesia. First, the skin above the skull over the telencephalon and the tectum was removed by using micro knives (10315-12, Fine Science Tools) (Figure S1A1). After drying the surface, the dental bond (Scotchbond Universal Adhesive, 41255, 3M) (shown in yellow) was pasted on the skull using a toothpick and illuminated it with blue LED light for 10 seconds (Figure S1A2). The u-shape metal was placed on the skull (Figure S1A3) and fluidic dental cement (Filtek Supreme Ultra, 6032XW, 3M) was placed on the skull (Figure S1A4, shown in white). To make the cement hard and fix the u-shape metal on the skull, blue LED light was illuminated for 10 seconds (Figure S1A4). The base of the fixation apparatus, the assembled harness, and the plastic ceiling to hold the fish body (Figure S1A5) were screwed (Figure S1A6). The tips of u-shape metal were inserted into the slits in the small part of assembled harness (Figures S1A7) and additional dental cement was placed on the contact point between them and illuminate them with the blue LED light for 10 seconds (Figure S1A8). After recovery from the anesthesia by perfusion with fish rearing water, the tethered fish with the fixation apparatus was transferred to the tank for the virtual reality environment (Figure S1A9). The tethered fish was kept at least 30 min in the dark condition under the microscope to habituate to the tethered situation with continuous perfusion of fish rearing water.

### GO/NOGO trial

Fish were trained for the GO trial (active avoidance) and the NOGO trial (passive avoidance) in the virtual reality environment under the microscope. Prior to the training, fish were kept separated in individual 1-litter tanks at least overnight. After the fixation to the custom-made harness followed by habituation to the tethered state, visual stimuli from 4 displays started to be presented and the adaptation session for visual stimuli was started. This adaptation session consisted of 20 GO/NOGO trials (10 GO trials and 10 NOGO trials with random order) without electric shock. After the adaptation session, we started the training session with electric shock. One training session consisted of 60 GO/NOGO trials. We mixed 10 GO and 10 NOGO trials in a random sequence in the first, middle and last 20 trials. Each fish performed 3-5 sessions. The inter-session interval was given for 15-20 minutes without showing visual stimuli. In the inter-trial interval, fish was shown white background with black stripes. In the GO trial, the background color in the vicinity of the fish in the virtual space changed to blue and that of the area ahead of the fish changed to red. In the NOGO trial, the color of the near side changed to red and that of the far side changed to blue. Success in the GO trial was defined by a correct escape to the red region by tail beats, which was initiated within 10 seconds after the change of the background color. Success in the NOGO trial was defined by a correct stay without tail beats in the red region for the 10 seconds after the change of the background color. Failure in the GO trial was a stay behavior without tail beats in the blue region after 10 seconds. Failure in the NOGO trial was an incorrect forward swim with tail beats into the blue region within 10 seconds. If the trials resulted in failure, electric shock (5V/cm for 1second) was delivered as a punishment from two needle electrodes put on both sides of the body of zebrafish (Figure 1A). In some fish, the color associated with the electric shock was reversed after the learning by the original rule was achieved. The other fish continued to perform by the original rule and, only if the second session showed learning behavior, the open-loop experiments (n = 11) and/or goal-color change experiments (n = 3) were applied.

In the open-loop experiment, we changed the arbitrary "gain" constant from 10 to 0. In this situation, fish tail beat frequency was multiplied with 0 and no feedback to virtual reality space was generated. In the goal color change experiment, we changed the goal background color from red to green or white as soon as fish reached the goal position.

### 2-photon imaging

To visualize calcium signals, we used 2-photon microscope LSM710-AX10 (Zeiss) with a water-immersion objective lens (W Plan-Apochromat, 20x, NA 1.0, Zeiss) and a mode-locked Ti: sapphire laser (Chameleon vision2, Coherent) at a wavelength of 890 nm. A 690-nm short-pass dichroic mirror (LP-690, Zeiss) was used to separate the excitation laser from the emitted fluorescence. Fluorescence emissions were collected using the GaAsP photomultiplier tube (BiG, Zeiss). The laser intensity was adjusted less than 150mW under the objective lens. The scanning was performed in bidirectional raster scanning with line step, 4. The speed of imaging is 97.75 ms per frame. The imaged field was 384.9 x 384.9 mm (512 x 512 pixels). Imaging was performed using a custom-made piezo actuator which allowed us to cover 3 planes separated by 16 μm.

### Analysis of neural activities

Image data analysis was performed using scripts written in ImageJ (ImageJ 1.50i and 1.51p, NIH), LabVIEW (National Instrument) and Matlab (Mathworks). The imaging data derived from one slice were collected using the custom macro of ImageJ. The collected image data of each slice were processed to remove the displacement of the x-y axis during the experiment. As reference image for such image registration, we used the average image of 1000 images with minimal displacement, from the dataset. We used 2-dimensional Fourier Transformation for images to find the displacement of individual images compared to the reference image in the frequency space. In case there was some degree of displacement in an image, with reference to the reference image, the distance or degree of displacement in the frequency domain was calculated in number of pixels along x and y axis by using cross-correlation. After calculating such relative displacement, each image was corrected for the shift in x and y axis. This method provided easier and more effective image registration compared to TurboReg plugin module for ImageJ. After the correction of the displacement, a median filter (radius, 2) was used to smoothen the image. Then the images were downsized from 512×512 to 256×256 to decrease the amount of further calculation time. In the next step, cell-like ROIs were detected from the image data. To define the ROIs corresponding to individual neurons, we used the following method. From the imaging data, peaky-ness of all pixels, as described in the method section of Ahrens et al., 2012 (Ahrens et al., 2012), was calculated. This peaky-ness corresponded to the change of activity of each pixel over time. Pixels inside a cell-body usually showed high peaky-ness value, as the activities of the cell caused a larger deviation of the pixels it includes, compared to pixels that do not belong to a cell. In the next step, we ranked the pixels with their peaky-ness values. A threshold peaky-ness value was calculated for later use by taking the average of these values. We took the pixel with the highest peaky-ness, and calculated the correlation of intensity along time between this pixel and surrounding pixels (51×51 pixels). We set a threshold value of correlation empirically, and if the correlation between this pixel and one neighboring pixel was higher than the threshold, that neighboring pixel was thought to be in the same cell as this pixel. By this process, a boundary of a cell-body could be obtained. We then looked for the pixel with the second-highest peaky-ness value in the updated list, and repeated the procedure described above to find another cell-body. This process was repeated to find cell bodies until the peaky-ness value of the pixel being considered became less than threshold peaky-ness value calculated earlier. After detection of potential cells in this method, we checked the spatial overlap between cells. If two cells had overlap, the one with smaller area was removed. Also, cells detected in the area of the image where there was apparently no brain tissue were also removed after careful visual observation. After these procedures, the fluorescence timeline of each cell was obtained. For each cell, fluorescence timelines of each pixel belonging to that cell were calculated from the image data, and an average timeline was calculated as the fluorescence of that cell. The fluorescence timelines for all cells from each slice/layer were calculated for each session of the experiment. As we recorded cellular activities from three layers, we gathered timelines of fluorescence for cells detected in all three layers. However, as each cell had only one intensity value in every three microscopic frames due to intra-layer switching, we used spline interpolation (built in function in LabVIEW) method to infer missing fluorescence values for each frame. With these, finally we could gather all the cells we could detect over three layers with equal length of intensity timelines. We calculated baseline intensity of every cell by averaging the baselines of their pixels, which were calculated by taking the average fluorescence intensity over the whole experiment time. To match the behavior data one to one to the fluorescence data, we checked the frame number of corresponding images taken by the microscope during the experiment, and cut the time-points from fluorescence data that did not correspond to behavior data. Finally, we calculated and calculate ΔF/F for each cell using the baseline defined above. By this method, we could observe the time-lapse change of activities of approximately 488 cells (n = 13) on the average in the three focal planes.

### Template matching

Template matching was performed as previously described (Bermudez Contreras et al., 2013) to evaluate whether specific neural activity patterns emerged during success trials in both GO and NOGO trials. Briefly, this method can quantify the similarity between a neuronal activity pattern (template) over a certain time window and other time windows of activity of the same length. In our case, we wanted to know if there were specific activity patterns of neurons that are responsible for the success of fish to avoid shock in a trial, and also whether the pattern emerges during the initial phase of a trial or not. For the creation of templates, we considered 10 GO and 10 NOGO trials that fish passed successfully (success rate > 80%) to reach the learner criteria, as the neural activity during this trials might have shown some specific pattern for success. We created a success GO template by taking the initial 2-seconds of neural activity for these 10 success GO trials at learner state, and taking an average over number of such trials. Similarly, we created a success NOGO template in the same manner. Then we took the neural activity data of the whole experiment, and calculated the similarity indices between the above mentioned templates and sliding time windows of 2-seconds length of the whole experimental data. After doing this, we could derive two graphs of similarity index, one for each template.

### Non-negative Matrix Factorization

To reveal further about encoded information in neurons, we performed Non-negative Matrix Factorization (NMF)(Figure 3A)(Lee and Seung, 2001). NMF separates synchronized neuronal activities, which enables the analysis of the neural ensembles. The data matrix D stores neuronal activities represented by their ΔF/F, where each column vector stores neuronal activities in each time frame. NMF searches for two factor matrices (P and T) such that matrix D can be best approximated by the product of matrices P and T by minimizing the error function: Error = ||*D* − *PT*||^2^. A gradient descent method was used to minimize the error function for each given matrix D. In each case, there are 5 independent attempts with randomly assigned P and T.

Maximum number of iteration for each attempt was set to 4000. The factorization minimizing the error function was taken to be the best factorization. The column vectors of the factor matrix P show typical neuronal activity patterns, *i.e.,* neural ensembles. The corresponding row vectors of the factor matrix T show the time course of patterns. The number of patterns, *i.e.,* the number of columns of the left factor matrix was determined by Akaike Information Criteria (AIC) (Figure 3A).

### Quantification of activities of neuronal ensemble

For the quantification of the change in the activity of the ensemble encoding the error from the prediction of the goal color, the peak values during first 10 GO trials, which is corresponding to the initial phase of learning, all success GO trials after establishment of learning and all GO trials in the open-loop trial were compared by the one-way ANOVA and Bonferroni's test. The change in the activity of the ensemble encoding the error from the prediction of the visual backward movement was also similarly compared among the initial phase of learning, in the success GO trials in the post-learning period, in the failure GO trials in the post-learning period, and in the open-loop trials. The activity of the ensemble encoding the error from the prediction of staying in the red goal region was also similarly compared between the state when fish was staying in the original red goal color and the states when the color of the goal was changed either to green or white.

