## Supplementary Information and Figures for "Future state prediction errors guide active avoidance behavior by adult zebrafish"

**Supplementary Figures**

**Figure S1. The fixation of living adult zebrafish, Related to Figure 1.**

(A) The procedure of the fixation of living adult zebrafish using the custom-made harness.

(A1) Scalp the skin above the telencephalon and the tectum. The skull was kept intact. (A2) After drying the surface, paste the dental bond (shown in yellow) using a toothpick and illuminate it with the blue LED light for 10 seconds. (A3) Place the u-shape metal on the skull. (A4) Put the dental cement (shown in white) to wrap around the u-shape metal using a toothpick and illuminate it with blue LED light for 10 seconds. (A5) Assemble the harness for the fixation. (A6) Place the fish on the base of the fixation apparatus and fix the harness to the base by two screws together with the plastic ceiling. (A7) Insert the tips of u-shape metal into the slits of the assembled harness. (A8) Put dental cement to the contact point between u-shape metal and the slits of harness and illuminate it with blue LED light for 10 seconds. (A9) Put the two needle electrodes for shock.

(B) The images of the custom-made harness. Left panel, disassembled; center panel, assembled; right panel, assembled with fish (nacre background).

**Figure S2. The activity of each neuron in the ensemble which emerged around the border into the goal, Related to Figure 3D.**

(A) The activity of each neuron averaged from 10 success GO trials after learning aligned according to the order of the timing of maximum activity.

(B) The activity of each neuron in 10 success GO trials (the same data in (A)).

The order of the sequential neuronal activation was largely kept in all 10 GO trials, suggesting the robustness of encoding position by each neuron in this ensemble.

**Movie S1, Related to Figure 1C**

The movies of the neural activities in the same frames as shown in the Figure 1C

### Figure S1

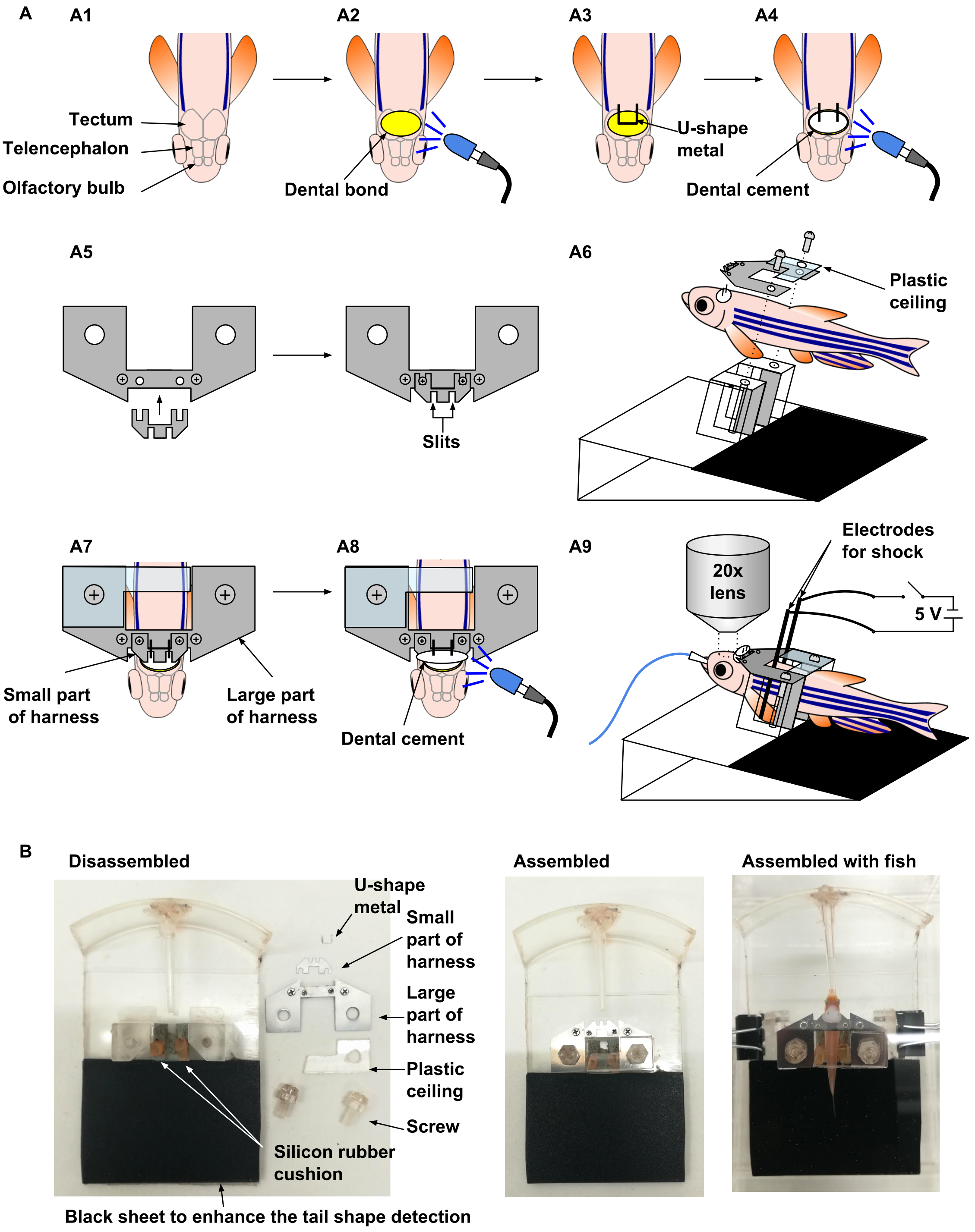
A
A1
A2
A3
A4
Tectum
Telencephalon
Olfactory bulb
Dental bond
U-shape
metal
Dental cement
A5
A6
Slits
Plastic
ceiling
A7
A8
A9
Small part
of harness
Large part
of harness
Dental cement
20x
lens
Electrodes
for shock
5 V
B
Disassembled
Assembled
Assembled with fish
U-shape
metal
Small
part of
harness
Large
part of
harness
Plastic
ceiling
Screw
Silicon rubber
cushion
Black sheet to enhance the tail shape detection

### Figure S2

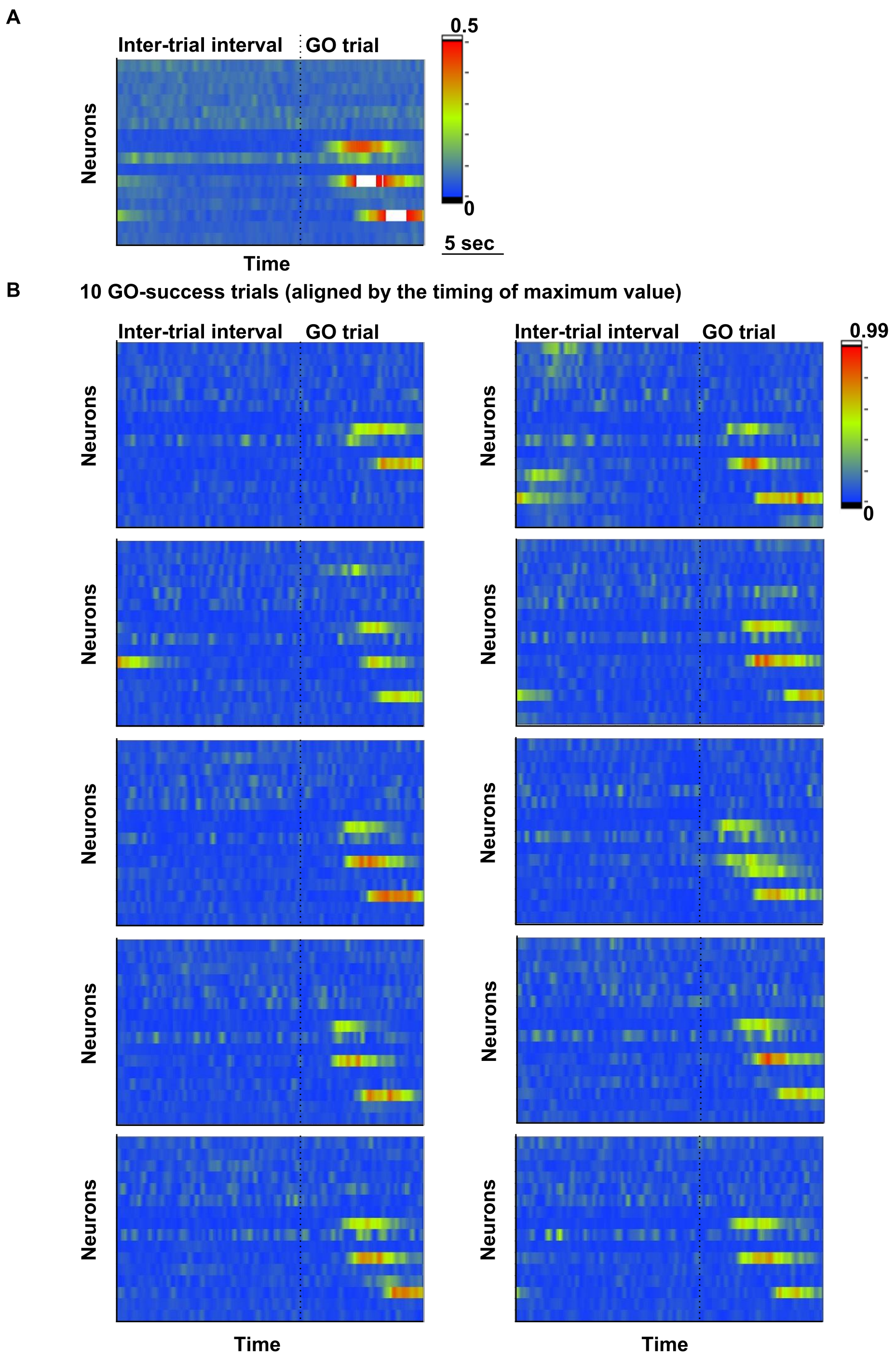
A
Inter-trial interval
GO trial
Neurons
Time
0.5
0
5 sec
B
10 GO-success trials (aligned by the timing of maximum value)
Inter-trial interval
GO trial
Neurons
0.99
0
Time
Time
